# Genotypic Detection of Antibiotic-Resistant Genes in Emerging Bacterial Pathogens From Neonatal Sepsis Patients in Lagos

**DOI:** 10.1101/2025.06.02.657359

**Authors:** Helen Opeoluwa Adegboyega, Taiwo Abayomi Banjo, Anotu Mopelola Deji-Agboola, Bamidele Abiodun Iwalokun, Wasiu Bamidele Mutiu, Olubunmi Adetokunbo Osinupebi, Tessie Owolabi Shorunmu, Farouk Adedeji Oladoja, Oluwabunmi Motunrayo Fatungase, Muinat Moronke Adeyanju, Raheem Akinwunmi Akindele, Ponle Bamidele Fakunle, Ayobami Babatunde, Olusola Ajayi, Olufemi Onadeko, Abimbola Adeola Akintunde Oyelekan, Solomon Olubodunrin Ariyibi, Kolawole Sunday Oritogun, Emmanuel Sunday Omirin

## Abstract

High-risk neonatal sepsis resulting in up to four million infants’ mortality and organ damages during the first 28 days of life predominates in low-income countries. We investigated genotypic antibiotic-resistant genes in emerging bacterial pathogens from newborn sepsis cases in Lagos.

A cross-sectional study of 294 at-risk hospitalised newborns in prominent paediatrics hospitals throughout three senatorial districts of Lagos utilised the socio-demographic data of their mothers and informed consents. We aseptically collected approximately 3 mL of venous blood from neonates, subsequently isolated and identified the associated bacteria using agar culture on Mueller Hinton agar. To evaluate the antibiotic resistance genes, we used Kirby-Bauer disc diffusion technique; MICROBACT^TM^ Identification System and molecular techniques including PCR, RAPD-PCR, sequencing and docking.

From 294 patient samples, 110 (37.4%) were cultured positive, with men (47%), females (53%), ages ≤ 72 hours (71.8%), and ages > 72 hours (28.1%). *Klebsiella pneumoniae, K. oxytoca, Enterobacter gergoviae, Serratia rubidiae, Escherichia coli,* and *Cronobacter sakazakii* were the involved Gram-negative bacteria, whereas Coagulase-negative *Staphylococci*, *Staphylococcus aureus*, and *Enterococcus faecalis* were the involved Gram-positive bacteria. Fourteen (12.7%) positive bacteria were resistant to only three antibiotic classes, whereas 73 (66.4%) were resistant to over three antibiotics. Antibiotic-resistant genes bla-CTX, bla-SHV, and bla-TEM were found in *Serratia rubidiae* and *Enterobacter gergoviae*, unusual aetiology of newborn sepsis. Molecular docking proved Meropenem’s antibacterial effectiveness.

The identified antimicrobial-resistant genes are crucial for improved patient management, as high-resistance strains of ampicillin, gentamicin, and cefotaxime remain ineffective against bacterial sepsis, leaving Meropenem as the sole feasible therapeutic option.

## INTRODUCTION

In the last two decades, improvements in child survival remains one of the major expectations in life expectancy at birth as the world’s neonatal mortality rate keeps falling from 37 deaths per 1,000 live births in 1990 to 17 per 1,000 live births in 2022. Approximately 4 million neonates die annually due to bacterial sepsis within the first 28 days of life, particularly in the low and middle-income countries with severe infection taking up to approximately 28% of the total deaths [1]

Neonatal sepsis (NS), systemic bacterial infections of newborn infants, is a major global public health challenge of infants linked to delicate neonatal period of life, either vertically transmitted “early-onset”-sepsis (EOS) within 72 hours of life; and/or horizontally transmitted “late-onset”- sepsis (LOS), after 72 hours of life. High-risk neonatal sepsis characterised by dysregulated host response to bacterial infection, is a leading predisposing factor to severe organ dysfunction, accounting for nearly a quarter of all newborn deaths.

At a global level, despite the seeming reduction in neonatal sepsis, meningitis, stillbirths and death occurring at the perinatal periods [2]; yet, the danger of antimicrobial resistance attributable to resistance to first-line antibiotics remains a threat to neonatal survival. Antibiotic resistance in hospital-acquired infections is developing due to poor infection control, novel microbial agents, and uncontrolled antibiotic prescription.

Could genotypic detection of newly emerging multiple drug resistance (MDR) bacterial genes be a significant risk factor for newborn sepsis? This three-year study aims to investigate the genotypic identification of antibiotic-resistant genes in emerging bacterial infections in newborn infants at three major government hospitals in Lagos, with the goal of assuring maternal-neonatal health and wellbeing in expansive enabling environments.

## MATERIALS AND METHODS

### SAMPLE COLLECTION, BACTERIA ISOLATION & IDENTIFICATION

We conducted a cross-sectional study of 294 at-risk newborns in three Lagos government pediatrics hospitals following patientś socio-demographic data, sought and obtained voluntary informed consents-to-participate in line with the Helsinki Declaration from mothers; having obtained Ethical approval (IBR/19/017) from Nigeria Institute of Medical Research (NIMR), Lagos.

We administered structured questionnaires in strict confidentiality, processing patient’s data (medical records, aged ≤ 28 days with fever (≥ 38°C), shock, other suspected classical symptoms of sepsis and positive blood culture)), while handling the babies’ clinical samples under the supervision of the paediatricians and human rights experts. Patients with congenital disorders and/or aged > 28 days were excluded from the study.

About 3mL of neonates’ venous blood were collected directly from each of the 294 patients into labelled pediatrics blood culture signal bottles under aseptic conditions were screen daily for evidence of bacterial growth within the period of incubation at 37°C for 1-7 days. Sequel to macroscopic evidence of no bacterial growth—absence of turbidity and gas formation, after 48 hours, the blood samples were sub-cultured onto Chocolate, Blood, MacConkey and Sabouraud agar plates, then incubated at 37 °C for 24 hours, targeting only aerobic organisms. Gram staining techniques were carried out on all isolated bacterial agents then identified by Agar culturing, and Mueller Hinton agar. Preservation and storage of all bacteria strains at -80 °C in Brain Heart Infusion (BHI) broth with 30 % glycerol until further tests.

We used the MICROBACT**^TM^** Identification System 12A, a qualitative micro-method based on enzyme reactions to identify aerobic and facultatively anaerobic Enterobacteriaceae, as well as other Gram-negative bacteria, using biochemical substrates and detection of colony features and biochemical properties (3). In addition, we adopted the Kirby-Bauer disc diffusion method on Mueller Hinton agar (Oxoid, U.K.) as recommended by the Clinical and Laboratory Standard Institute (4)

Eighteen different antibiotics examined included ampicillin (10 µg), oxacillin (1µg), amoxicillin–clavulanic acid (30 µg), cefoxitin (30 µg), cefotaxime (30 µg), ceftriaxone (30 µg), ceftazidime (30 µg), cefuroxime (30 µg). Others were meropenem (10µg), tazobactam, (110µg), levofloxacin (5 µg), clindamycin (2µg), cefepime (30µg), vancomycin (10 µg), gentamycin (10 µg), amikacin (30 µg), erythromycin (15 µg) and ciprofloxacin (5 µg). The plates were incubated at 37^0^C for 24 hours while the diameter of zones of inhibition of each antibiotic measured in millimeters against the bacteria were documented and interpreted appropriately (5).

All tests were controlled with standard organisms: *E. coli* (ATCC 35218), *E. coli* (ATCC 25922) for the Gram-negative isolates; and *Staphylococcus aureus* (ATCC 29213) for Gram-positive isolates. Appropriate biochemical tests were performed on all isolates and the standard organisms; while further tests against the same batch of antibiotics using plain Mueller Hinton were carried out.

The beta-lactamases activity was determined using a chromogenic cephalosporin method (**6**) (Nitrocephin-stick, oxoid, UK), using *Staphylococcus aureus* (ATCC 29213) as positive control. Bacterial isolates resistant to third generation cephalosporins: ceftazidime, cefotaxime and ceftriazone were tested for extended spectrum beta lactamases (ESBL) production by Double-disk synergy test (DDST) to detect ESBL producers. Cefotaxime-clavulanic disk was placed 20 mm apart center to center.

## MOLECULAR ISOLATION & IDENTIFICATION OF GENOMIC DNA

### Molecular Characterization of Isolates

To establish the molecular characteristics of the bacterial isolates, using PCR, RAPD-PCR, gene sequencing and docking techniques, bacterial DNA isolation from the samples were carried out in a stepwise procedure using the alkaline lysis (spin column) method according to the NIMR Biotech DNA Extraction Kit Protocol according to the protocol of Trindade et al. (7)

DNA was collected by centrifugation at 13,000 x g for 10 min, washed with 500 μl of 70 % ethanol, air-dried at room temperature for approximately three hours and finally dissolved in 50 μl of Tris-EDTA (TE) buffer. The quality of the extracted DNA was measured 1.7-2.0 using nano-drop spectrophotometer (Thermo Fisher Scientific, Wilmington, Delaware, USA) and the absorbance recorded at a wavelength of 260/280 nm and concentration not less than 5.0 ng/µl. Pure DNA was isolated, stored and sealed at -20 °C for future.

Polymerase Chain Reaction (PCR) were performed using Pelter thermal cycler (MJ Research, BIORAD Alpha unit block assembly for DNA engine systems. U.S.A.) in a 25 µl reaction mixture using gene specific primers according to the methods of Kramer and Coen. (**8**)

To detect the presence of the ESBL enzyme in the bacteria, a set of primers were used: Solis Biodyne 5x HOT FIRE Pol Blend Master mix in a multiplex Polymerase chain reaction:

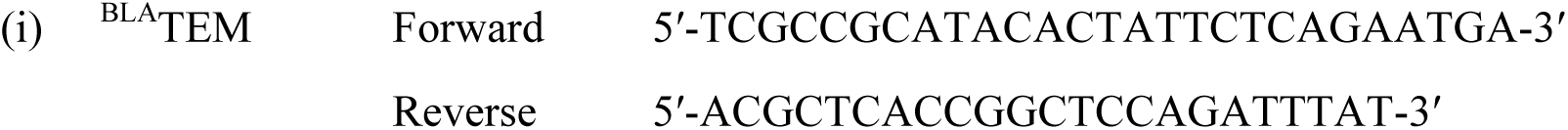
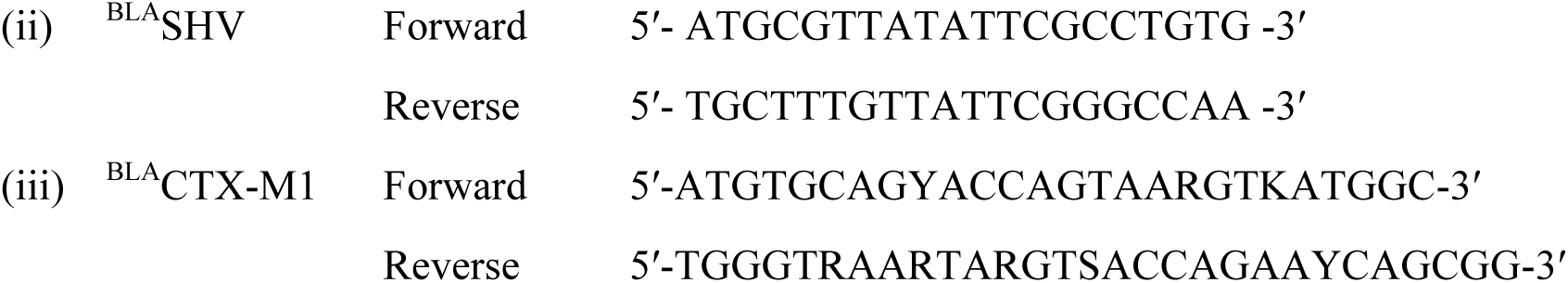

Thermo-Scientific GeneRuler 100 bp DNA ladder was used as the DNA molecular weight standard, the DNA was characterised using electrophoresis followed by visual assessment of the bands by ethidium bromide dye aided UV-Trans-illuminator. Subsequently, amplified fragments were sequenced using a Genetic Analyzer 3130 xl sequencer (Applied Biosystems), sequencing kit (Big Dye terminator v3.1 cycle), Bio-Edit software and MEGA 6 for all genetic analysis-7 afterwards, molecular docking.

### Statistical analysis

Data acquired in this study were analyzed using Microsoft Excel, Epi-Info 6.0, and Graph Pad Prism. The treatment group data, presented as mean ± SEM, were analyzed using T-test and chi- square, followed by SPSS analysis to ascertain the association between variables and their significance levels (p < 0.05).

### Ethical considerations

Ethical approval and permission were obtained from the Institutional Research Board, Nigeria Institute of Medical Research, Lagos, following adherence to their existing bioethical regulations. Voluntary informed consent was solicited and acquired from all parents and minors’ guardians to participate, while clarifying the study’s objectives, advantages, and confidentiality of the information submitted. The research was conducted in line with the Helsinki Declaration.

This article does not disclose personal data that could identify patients, as we strictly complied with patient confidentiality and privacy rights in accordance with the approved publication process.

## RESULTS

Findings from this study revealed One hundred and ten (37.4 %) of the initial 294 blood samples collected were positive cultures. Seventy-nine (71.8%) and 31 (28.1 %) were aged ≤ 72 hours and > 72 hours respectively; presented as 52 (47 %) males and 58 (53 %) females. **Table 1** gives the detailed information of both the maternal (gestational age, pre-delivery conditions, mode of deliveries) and neonatal (age at term, birth weight and age at onset of bacterial sepsis) demographics.

**TABLE 1:** DEMOGRAPHIC CHARACTERIZATION OF NEONATES WITH BACTERIAL SEPSIS.

| CHARACTERISTICS |  | n (%) |
| --- | --- | --- |
| **MATERNAL DATA** |  |  |
| **Gestational age** | Pre-term (<37 weeks) | 76 (69) |
| | Term (≥ 37 weeks) | 34 (31) |
| **Pre-delivery conditions** | Premature rupture of membrane | 28 (25) |
|  | Intrauterine distress | 15 (14) |
| **Mode of delivery** | Caesarian section | 65 (59) |
|  | Vagina | 45 (41) |
| **NEONATAL DATA** |  |  |
| **Gender** | Female | 58 (53) |
|  | Male | 52 (47) |
| **Age at onset of bacterial sepsis** | Early-onset (≤72 hrs) | 79 (72) |
|  | Late-onset (>72 hrs – 28 days) | 31 (28) |
| **Birth weight** | Very low birth weight (≤1500g) | 23 (21) |
|  | Low birth weight (> 1500g – 2500g) | 53 (48) |
|  | Normal (> 2500g) | 34 (31) |

*Term gestational age were 0.115 times less likely to have sepsis than preterm ( $p>0.05$ ). Normal birth weight were 0.929 times less likely to have sepsis than low birth weight ( $p>0.05$ ). Premature rupture of membrane were 1.296 times more likely to experience sepsis than those without PROM ( $p>0.05$ ).*

Out of 110 neonates with positive blood cultures, only 70 (64 %) grew Gram-negative bacteria while 40 (36 %) grew Gram-positive bacteria. The pathogens implicated in NS were indicated in **Table 2**.

**TABLE 2:** ANNUAL TREND OF BACTERIAL AGENTS IMPLICATED IN NEONATAL SEPSIS.

| **PATHOGENS** | **Year 1** | **Year 2** | **Year 3** | **Total (n, %)** | **Mean value** |
| --- | --- | --- | --- | --- | --- |
| *K. pneumonia* | 6 | 11 | 10 | 27 (24.5) | 9 |
| Coagulase negative staphylococci | 7 | 9 | 5 | 21 (19) | 7 |
| *Klebsella oxyoca* | 3 | 9 | 5 | 17 (15.4) | 5.7 |
| *Staphylococcus aureus* | 6 | 5 | 4 | 15 (13.6) | 5 |
| *Enterobacter gergoviae* | 3 | 5 | 2 | 10 (9.1) | 2 |
| * *Serratia rubidiae* | 4 | 3 | - | 7 (6.4) | 2.3 |
| *Escherichia coli* | 3 | 1 | 1 | 5 (4.5) | 1.7 |
| ** *Cronobacter sakazakii* | - | 2 | 2 | 4 (3.6) | 1.3 |
| *Enterococcus faecalis* | 1 | 2 | 1 | 4 (3.6) | 1.3 |
| **Total** | **33** | **47** | **30** | **110** |  |

Over the three-year study period, the four most prevalent pathogens namely *Klebsiella pneumoniae* 27 (24.5%), *Coagulase-negative Staphylococcus* 21 (19%), *Klebsiella oxytoca* 17 (15.5%) and *Staphylococcus aureus* 15 (14%), altogether accounted for 73 % of all confirmed cases of neonatal sepsis in the Lagos communities **(Table 2).** However, two newly identified isolates: *Serretia rubidiae* and *Enterobacter gergoviae,* seldomly known aetiological agents of neonatal sepsis were documented. Antibiotic resistance genes (ARG) encoding bla-CTX, bla-SHV, bla- TEM were detected by PCR molecular-typing techniques and their relatedness was captured by the phylogenetic tree.

Variation exists in the distribution of pathogens between the preterm and term infants according to their modes of deliveries. Preterm neonates (gestational age ≤ 37 weeks) presented high positive culture 76 (69 %) characterized by *Klebsiella pneumoniae* 20 (26.3 %), *Co-NS* 15 (19.7 %), *Klebsiella oxytoca* 10 (13.2 %) and *Staphylococcus aureus* 9 (11.8 %) unlike term-infants.

The infected term-infants (gestational age ≥ 37 weeks) with 34 (31 %) positive culture were characterized by *Klebsiella pneumoniae* 7 (20.6%), *Klebsiella oxytoca* 7 (20.6 %), *Co-NS* 6 (17.6%) and *Staphylococcus aureus* 6 (17.6%). Meanwhile, infants’ deliveries with history of PROM lacked *C. sakazakii* and *E. faecalis* infections while deliveries with intrauterine distress (IUD) had no exposure to *S. rubidae* infection **(Table 3)**

**Table 3:**
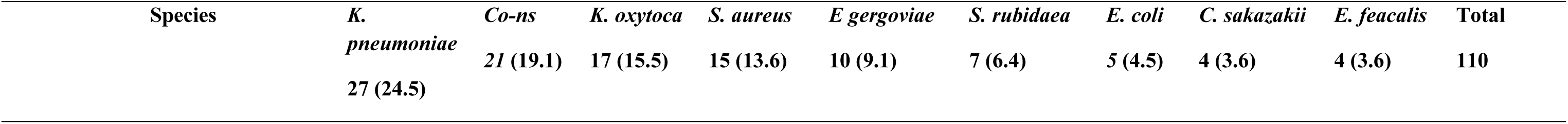
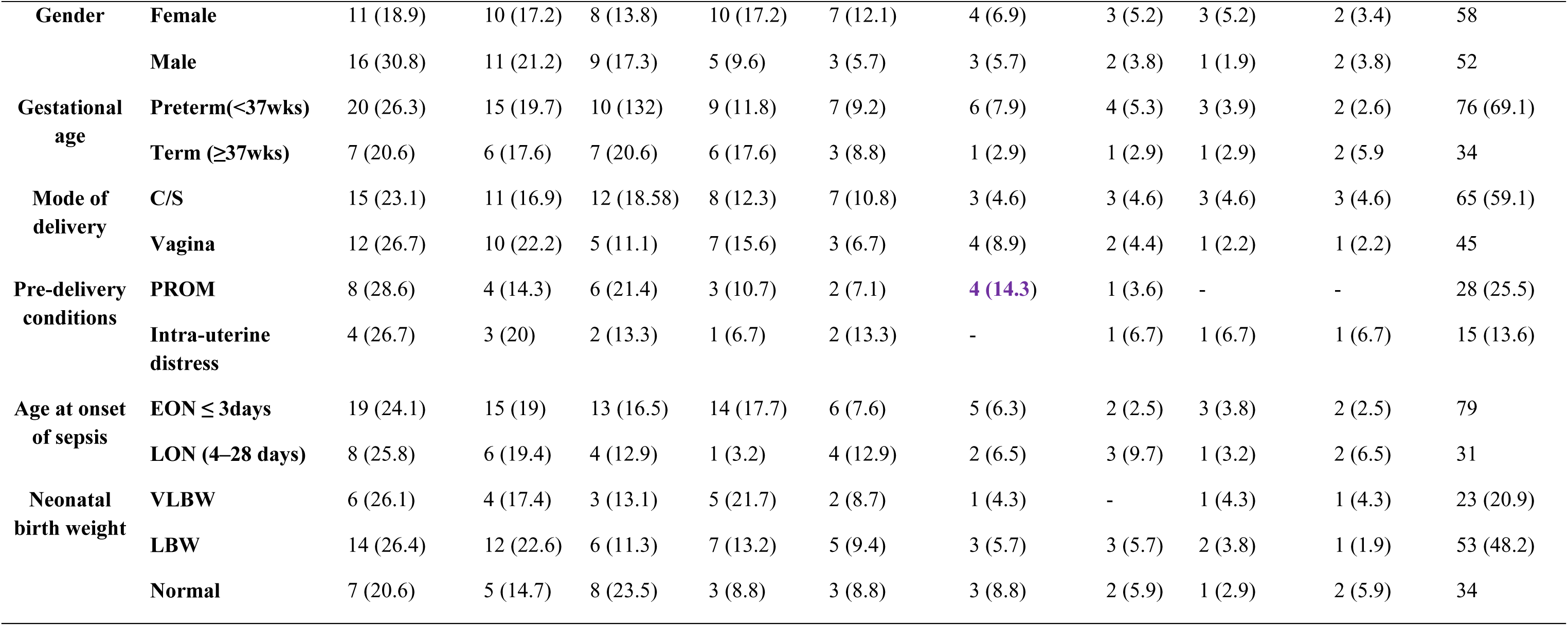
BACTERIAL AGENTS & RISK FACTORS ASSOCIATED WITH NEONATAL SEPSIS.

| Species | *K. pneumoniae* | *Co-ns* | *K. oxytoca* | *S. aureus* | *E gergoviae* | *S. rubidaea* | *E. coli* | *C. sakazakii* | *E. feacalis* | Total |
| --- | --- | --- | --- | --- | --- | --- | --- | --- | --- | --- |
|  | 27 (24.5) | 21 (19.1) | 17 (15.5) | 15 (13.6) | 10 (9.1) | 7 (6.4) | 5 (4.5) | 4 (3.6) | 4 (3.6) | 110 |

|  |  |  |  |  |  |  |  |  |  |  |  |
| --- | --- | --- | --- | --- | --- | --- | --- | --- | --- | --- | --- |
| **Gender** | **Female** | 11 (18.9) | 10 (17.2) | 8 (13.8) | 10 (17.2) | 7 (12.1) | 4 (6.9) | 3 (5.2) | 3 (5.2) | 2 (3.4) | 58 |
|  | **Male** | 16 (30.8) | 11 (21.2) | 9 (17.3) | 5 (9.6) | 3 (5.7) | 3 (5.7) | 2 (3.8) | 1 (1.9) | 2 (3.8) | 52 |
| **Gestational age** | **Preterm(<37wks)** | 20 (26.3) | 15 (19.7) | 10 (13.2) | 9 (11.8) | 7 (9.2) | 6 (7.9) | 4 (5.3) | 3 (3.9) | 2 (2.6) | 76 (69.1) |
|  | **Term (≥37wks)** | 7 (20.6) | 6 (17.6) | 7 (20.6) | 6 (17.6) | 3 (8.8) | 1 (2.9) | 1 (2.9) | 1 (2.9) | 2 (5.9) | 34 |
| **Mode of delivery** | **C/S** | 15 (23.1) | 11 (16.9) | 12 (18.58) | 8 (12.3) | 7 (10.8) | 3 (4.6) | 3 (4.6) | 3 (4.6) | 3 (4.6) | 65 (59.1) |
|  | **Vagina** | 12 (26.7) | 10 (22.2) | 5 (11.1) | 7 (15.6) | 3 (6.7) | 4 (8.9) | 2 (4.4) | 1 (2.2) | 1 (2.2) | 45 |
| **Pre-delivery conditions** | **PROM** | 8 (28.6) | 4 (14.3) | 6 (21.4) | 3 (10.7) | 2 (7.1) | 4 (14.3) | 1 (3.6) | - | - | 28 (25.5) |
|  | **Intra-uterine distress** | 4 (26.7) | 3 (20) | 2 (13.3) | 1 (6.7) | 2 (13.3) | - | 1 (6.7) | 1 (6.7) | 1 (6.7) | 15 (13.6) |
| **Age at onset of sepsis** | **EON ≤ 3days** | 19 (24.1) | 15 (19) | 13 (16.5) | 14 (17.7) | 6 (7.6) | 5 (6.3) | 2 (2.5) | 3 (3.8) | 2 (2.5) | 79 |
|  | **LON (4–28 days)** | 8 (25.8) | 6 (19.4) | 4 (12.9) | 1 (3.2) | 4 (12.9) | 2 (6.5) | 3 (9.7) | 1 (3.2) | 2 (6.5) | 31 |
| **Neonatal birth weight** | **VLBW** | 6 (26.1) | 4 (17.4) | 3 (13.1) | 5 (21.7) | 2 (8.7) | 1 (4.3) | - | 1 (4.3) | 1 (4.3) | 23 (20.9) |
|  | **LBW** | 14 (26.4) | 12 (22.6) | 6 (11.3) | 7 (13.2) | 5 (9.4) | 3 (5.7) | 3 (5.7) | 2 (3.8) | 1 (1.9) | 53 (48.2) |
|  | **Normal** | 7 (20.6) | 5 (14.7) | 8 (23.5) | 3 (8.8) | 3 (8.8) | 3 (8.8) | 2 (5.9) | 1 (2.9) | 2 (5.9) | 34 |

**TABLE 4:** MULTIDRUG RESISTANCE PATTERN OF BACTERIAL ISOLATES IN NEONATAL SEPSIS.

| Bacterial isolates | (N) | Degree of resistance | | | | | | | | | Total MDR isolated ≥ R3 |
| --- | --- | --- | --- | --- | --- | --- | --- | --- | --- | --- | --- |
|  |  | RO (%) | R1 (%) | R2 (%) | R3 (%) | R4 (%) | R5 (%) | R6 (%) | R7 (%) | R8 (%) | %MDR |
| *K. pneumoniae* | 27 | – | 3 (11.1) | 1 (3.7) | 6 (22.2) | 7 (25.9) | 10 (37.0) | – | – | – | 23 (85.2) |
| *CO-NS* | 21 | – | – | 5 (23.8) | 2 (9.5) | 1 (4.8) | 5 (25.8) | 2 (9.5) | 3 (14.3) | 3(14.3) | 16 (76.2) |
| *K. oxytoca* | 17 | 2 (11.8) | – | 1 (5.9) | 1 (5.9) | 7 (41.2) | 6 (35.3) | – | – | – | 14 (82.4) |
| *E. gergoviae* | 10 | – | – | -- | 1 (10) | 5 (50) | 4 (40) | – | – | – | 10 (100) |
| *S. aureus* | 15 | 1 (6.7) | 2 (13.3) | 3 (20) | 2 (13.3) | 3 (20) | 2 (13.3) | 1 (6.7) | - | 1 (6.7) | 9 (60) |
| *S. rubidiae* | 7 | – | – | – | 1 (14.3) | 2 (28.6) | 4 (57.1) | – | – | – | 7 (100) |
| *E. coli* | 5 | – | – | 2 (40) | – | 1 (20) | 2 (40) | – | – | – | 3 (60) |
| *C. sakazakii* | 4 | – | 2 (50) | 1 (25) | 1 (25) | – | – | – | – | – | 1 (25) |
| *E. faecalis* | 4 | – | – | – | – | – | – | – | 1 (25) | 3 (75) | 4 (100) |
| **Total** | **110** | **3** | **7** | **13** | **14** | **26** | **33** | **3** | **4** | **7** | **87** |

**TABLE 5:**
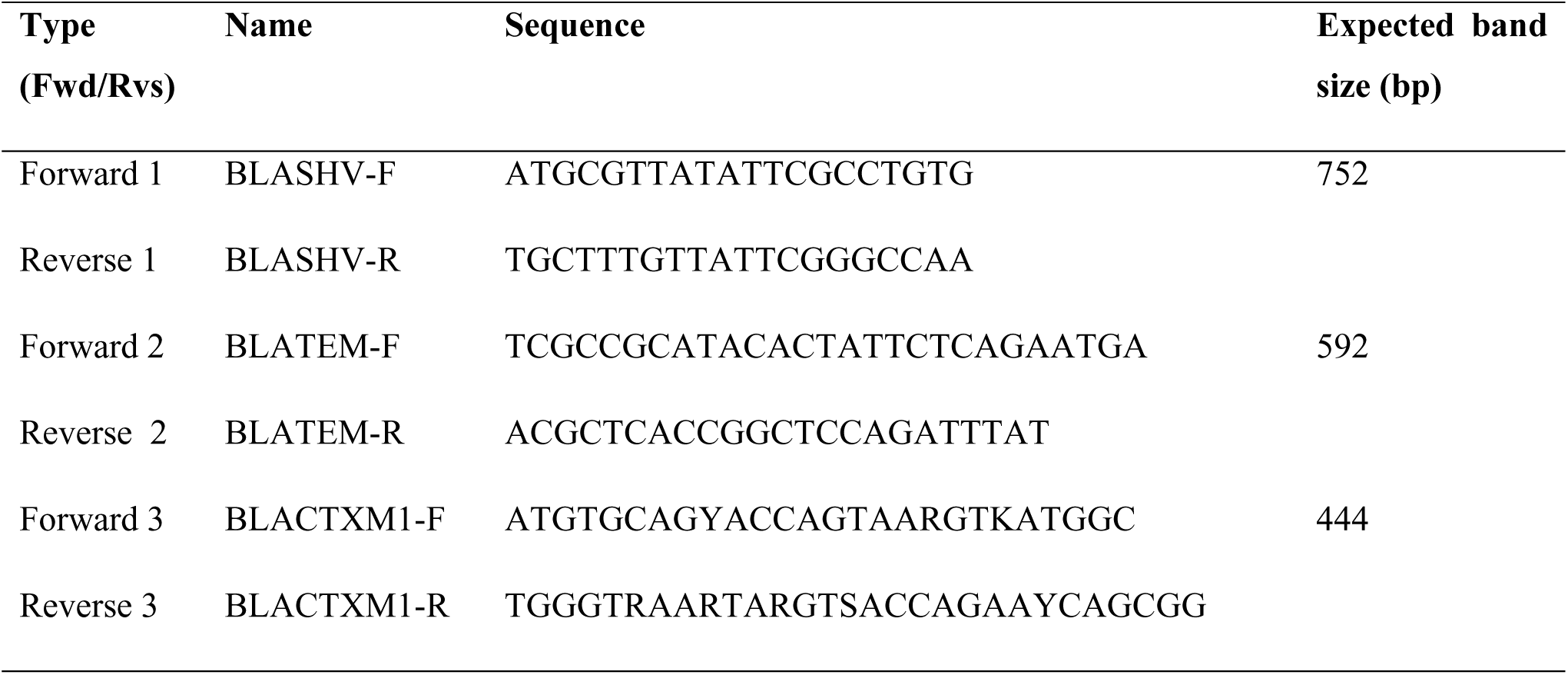
PRIMERS FOR THE AMPLIFICATION OF ESBL.

**TABLE 6:** PERCENTAGE DISTRIBUTIONS OF ESBLS GENES.

| GENE | FREQUENCY (#) | (%) | % MDR |
| --- | --- | --- | --- |
| SHV | 18 | 34.0 | 28.6 |
| CTX-M | 52 | 98.1 | 82.5 |
| TEM | 42 | 79.0 | 66.7 |
| CTX-M, TEM | 23 | 43.4 | 36.5 |
| TEM, SHV | 0 | 0 | 0 |
| CTX-M, SHV | 0 | 0 | 0 |
| CTX-M, SHV, TEM | 18 | 34 | 28.6 |

Preceding sequencing (**Figure 1**, **Table 7)** ten different bacterial isolates - five of *Enterobacter gergoviae* (lanes 69, 5, 50, 61, 51) and five of *Serratia rubidiae* (lanes 16, 3, 62, 54, 30) — previously exhibiting susceptibility to Meropenem, Imipenem (9) Ciprofloxacin, and Sulfamethoxazole (10) respectively displayed divergent sequence identities and concomitant antibiotic therapeutics.

**Figure 1:**
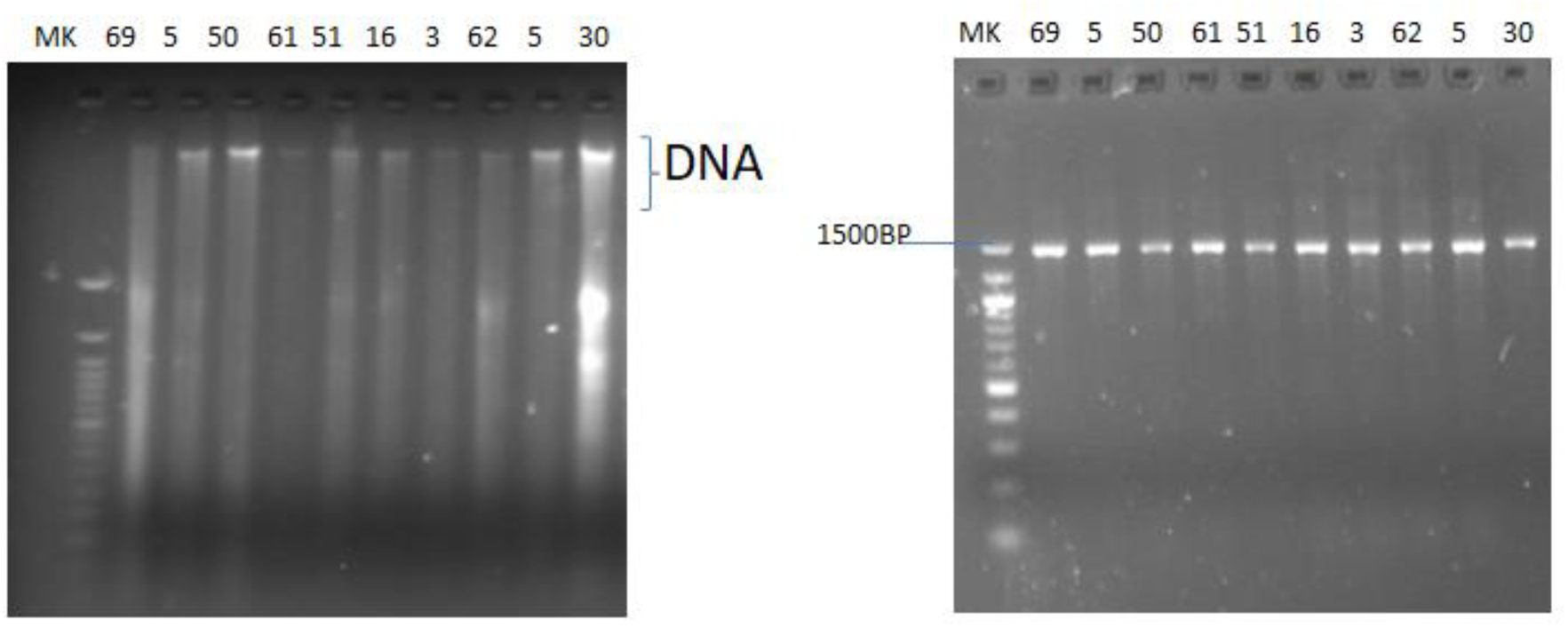
**(a)** Agarose gel electrophoresis of the DNA EXTRACTED from selected bacteria isolates. **(b)** Agarose gel electrophoresis of the PCR products 16S rRNA amplified from selected bacteria isolates. (Band size approximately 1500 bp). Gel image indicates a positive amplification in all samples

**TABLE 7:**
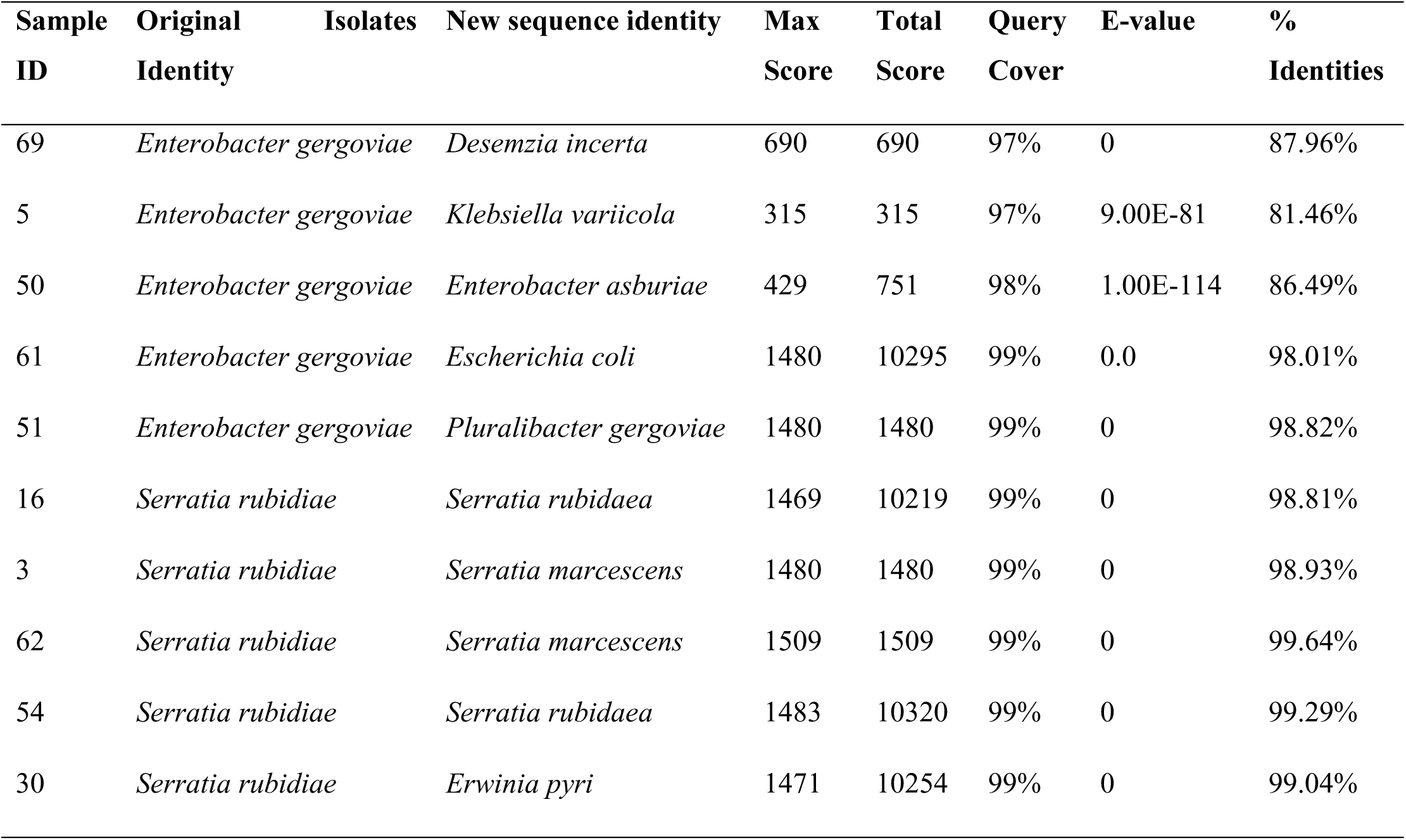
NCBI BLAST SEQUENCE CONFERRED DIVERGENT IDENTITY TO ORIGINAL BACTERIAL ISOLATES.

**TABLE 8:** MOLECULAR DOCKING OF DNA GYRASE (PDB ID: 6YD9)

| Ligands | Binding Energy<br>(kcal/mol) | Hydrogen Bond<br>Interaction |  | Hydrophobic<br>Interaction | Pi-interaction |
| --- | --- | --- | --- | --- | --- |
|  |  | Amino acid<br>residue | Distance<br>(Å) |  |  |
| **Piperacillin** | -7.9 | Asn46 | 3.05 | Val43, Ala47,<br>Val71, Ile94 | Val43, Ala47, Val71,<br>Ile94 |
| **Ciprofloxacin** | -7.8 | Gly77<br>Thr165 | 2.34<br>2.41 | Ala47, Ile78 | Ile78 |
| **Amoxicillin** | -7.2 | Asp73<br>Gly77<br>Val97<br>Thr165 | 2.11<br>2.71<br>2.84<br>2.90 | Ile94 | Ile94 |
| **Imipenem** | -7.0 | Glu50<br>Val71<br>Asp73<br>Arg76<br>Gly77<br>Thr165 | 3.10<br>2.12<br>1.98<br>2.48<br>2.66<br>3.08<br>2.60<br>2.69 | - | - |
| **Cefepime** | -6.9 | Asn46<br>Asp73<br>Thr165 | 3.09<br>1.95<br>2.78 | Val43, Asn46,<br>Ala47, Val71,<br>Thr165, Val167 | Val43, Ala47, Val71,<br>Thr165, Val167 |

**Figure 2:**
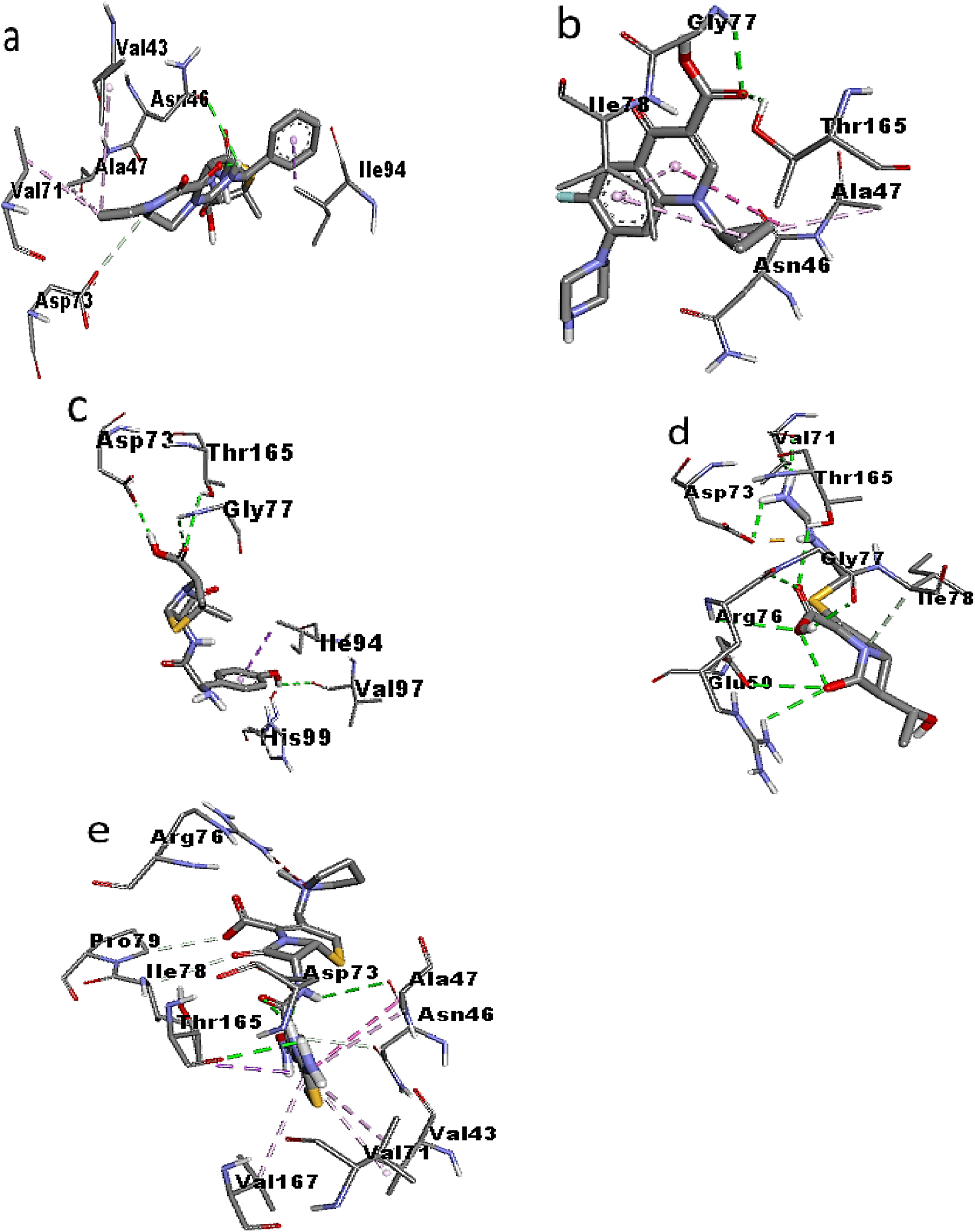
**Interaction diagram of 6YD9 with (a)** piperacillin, (**b)** ciprofloxacin, (**c)** amoxicillin, (**d)** imipenem, and (**e)** cefepime. **Keys***: Ala = Alanine, Arg =Arginine, Asn =Asparagine, Asp =Aspartic acid, Gln =Glutamine,* Glu= Glutamate, Gly= Glycine, *Ile =Isoleucine, Pro =Proline, Thr =Threonine, Val =Valine)*

**TABLE 9:**
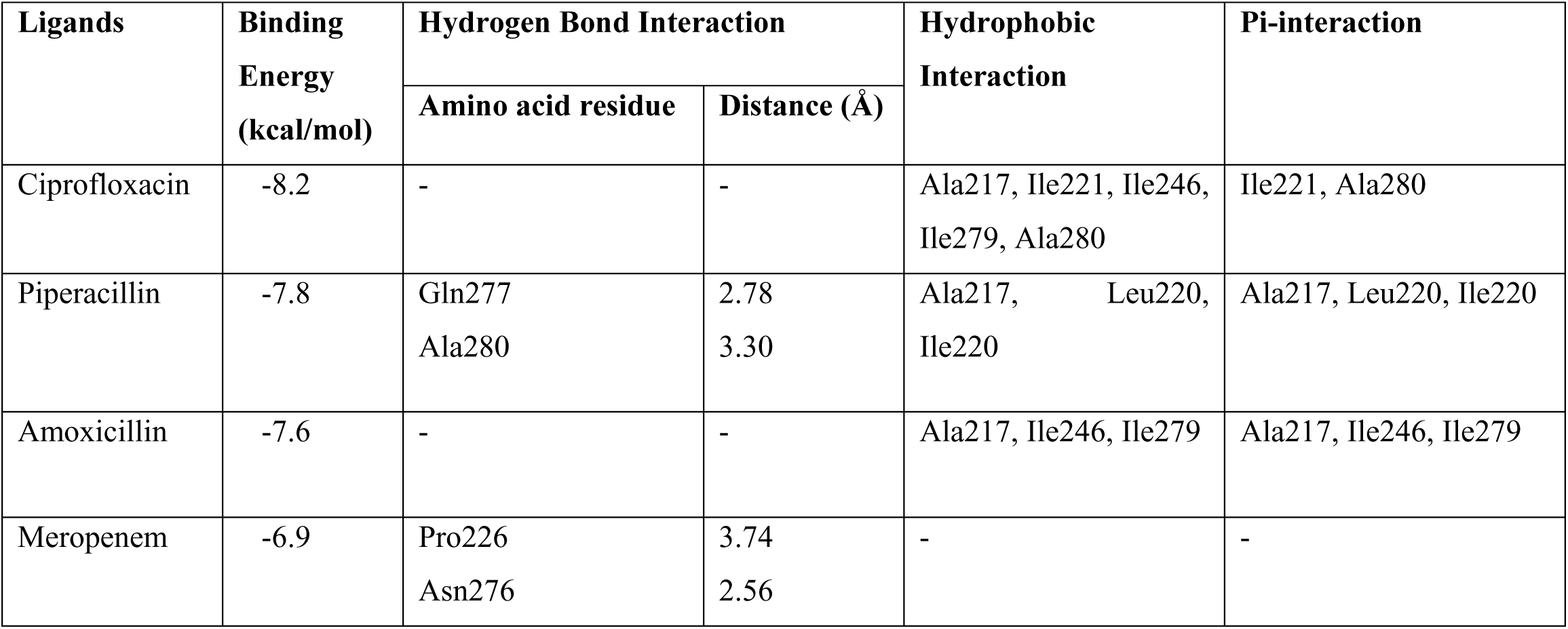
MOLECULAR DOCKING OF BETA LACTAMASE (PDB ID: 1SHV)

| Ligands | Binding Energy (kcal/mol) | Hydrogen Bond Interaction |  | Hydrophobic Interaction | Pi-interaction |
| --- | --- | --- | --- | --- | --- |
|  |  | Amino acid residue | Distance (Å) |  |  |
| Ciprofloxacin | -8.2 | - | - | Ala217, Ile221, Ile246, Ile279, Ala280 | Ile221, Ala280 |
| Piperacillin | -7.8 | Gln277<br>Ala280 | 2.78<br>3.30 | Ala217, Leu220, Ile220 | Ala217, Leu220, Ile220 |
| Amoxicillin | -7.6 | - | - | Ala217, Ile246, Ile279 | Ala217, Ile246, Ile279 |
| Meropenem | -6.9 | Pro226<br>Asn276 | 3.74<br>2.56 | - | - |

Sequenced *Enterobacter gergoviae* revealed *Desemzia incerta, Klebsiella variicola, Enterobacter asburiae, Escherichia coli, and Pluralibacter gergovaie, and* necessitated reconsideration of new antibiotics regime–Piperacillin-Tazobactam, Ciprofloxacin, Meropenem and/or Imipenem (11)); Amoxicillin/Clavulanate (12–14).

Meanwhile, the new sequence identities of *Serratia rubidiae were* susceptible to the following antibiotic regimes: Ciprofloxacin and Sulfamethoxazole (10), Cefepime (15) *Piperacillin- tazobactam,* Cefepime (16–15)

These trends in the comparative therapeutics of antibiotics, informed by recent sequencing evaluations, elucidate a primary factor contributing to antibiotic resistance genes and the potential for enduring neonatal sepsis when antibiotic selection relies solely on sensitivity analysis. for selecting antibiotic therapy.

**Figure 3:**
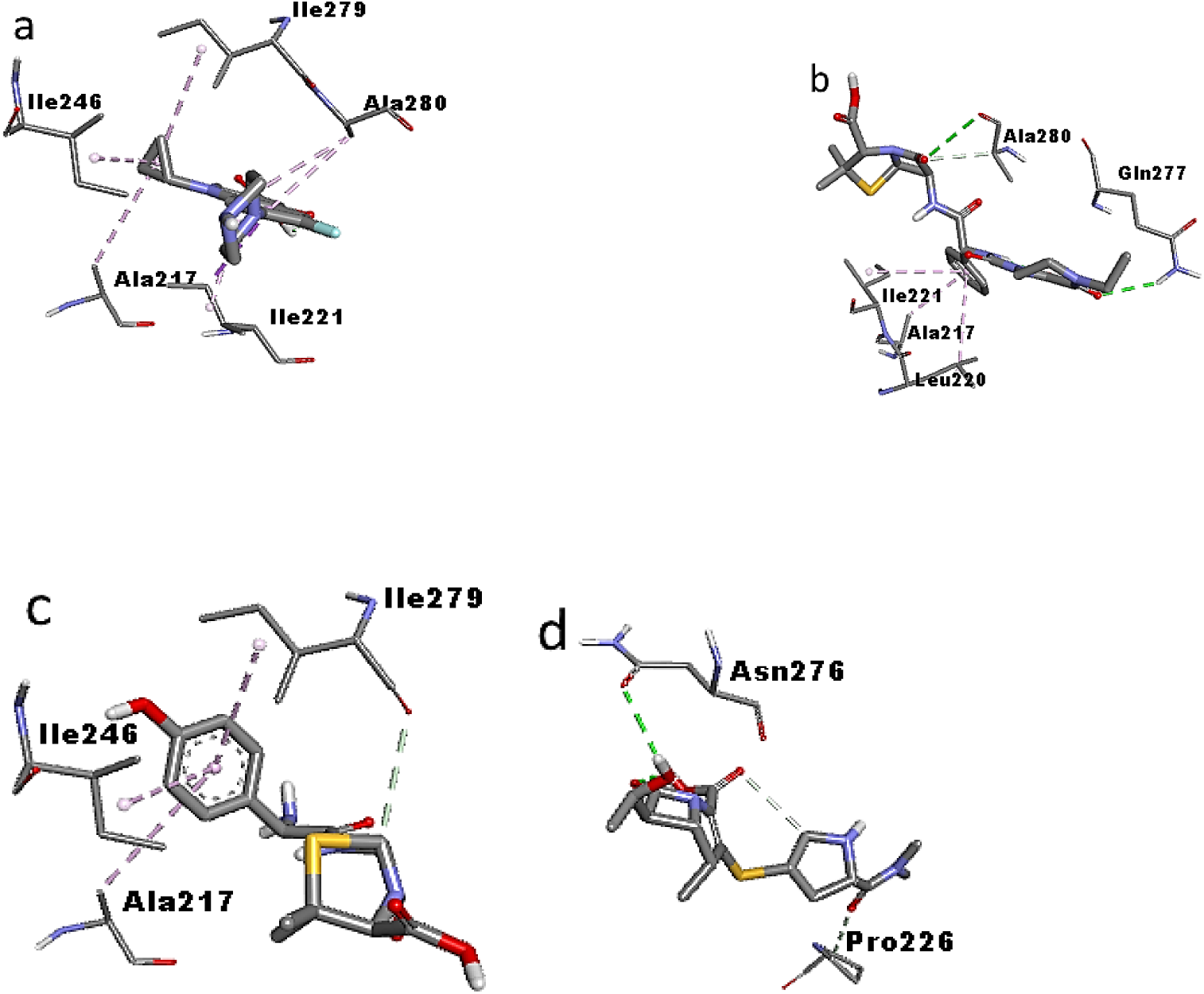
**Interaction diagram of 1SHV with (a)** ciprofloxacin, **(b)** piperacillin, **(c)** amoxicillin, **(d)** meropenem *(most preferred antibiotics for NS).* **Keys:** Ala =Alanine, Asn =Asparagine, Gln =Glutamine, Ile =Isoleucine, Pro = Proline.

## DISCUSSION

Neonatal sepsis (NS) remains a significant cause of neonatal morbidity posing a considerable health concern globally [1,17]. In the present study, 110 (37.4 %) of 294 high-risk neonates initially suspected of sepsis, were confirmed by positive culture. In 71.8 % of EOS cases, the predominantly isolated organism in descending order included *Klebsiella pneumoniae* (24.5 %), Coagulase-Negative Staphylococci (19.1 %), *Klebsiella oxytoca* (15.5 %), *Staphylococcus aureus* (13.6 %), *Enterobacter gergoviae* (9.1 %), and the least frequently identified pathogens were *Cronobacter sakazakii* and *Enterococcus faecalis* (3.6 %).

Multiple risk factors associated with NS in this cohort **(Table 3)** were prematurity (gestational age <37 weeks), low birth weight (LBW and very low birth weight, VLBW), intrauterine device (IUD, 14 %), and premature rupture of membranes (PROM, 25 %) (18–21) Similarly, prematurity (69.1 %) was a major vulnerability factor, reinforcing previously documented evidence of preterm infants being more susceptible to sepsis complications following underdeveloped humoral and cellular immunity (19, 20, 22).

The frequently implicated Gram-negative organisms in EOS demonstrated profound resistance against first-line antibiotics such as ampicillin and gentamicin, as well as cefotaxime, a common second-line agent, corroborating similar reports from other developing nations. In particular, *Klebsiella pneumoniae* has 96 % resistance to ampicillin, 100 % to gentamicin, and significant resistance to cefotaxime 85 %; Other Gram-negative isolates, including *Klebsiella oxytoca, Enterobacter gergoviae*, and *Serratia rubidaea,* and Gram-positive organisms like Coagulase-negative Staphylococci and Staphylococcus aureus have all replicated peculiar resistance profiles, supporting earlier findings by Rahman *et al*. [23).

The resistance landscape was particularly worrisome for aminoglycosides and β-lactam agents. Many Gram-positive isolates demonstrated high-level resistance to amikacin (67 – 100 %) and moderate-to-high resistance to gentamicin (53 –100 %). resistance levels to Amoxyclav (33 – 100 %).

Notably, all Gram-negative organisms exhibited complete resistance (100 %) to cefuroxime and very high resistance (>85 %) to a broad range of cephalosporins, including cefoxitin, ceftriaxone, ceftazidime, cefotaxime, and cefepime; resistance to Amoxyclav (50 – 100 %). Although the incidence of NS due to Enterobacteriaceae has declined to approximately 20% due to improved prenatal screening and intrapartum antibiotic administration [(24, 25–27) *Klebsiella spp., Enterobacter spp*., and *Serratia spp*. remain the principal aetiological agents in many cases (28). All these findings highlight the increasing prevalence of multidrug-resistant (MDR) organisms, which poses a substantial challenge to effective empirical antibiotic therapy.

Maternal risk factors– including prolonged rupture of membranes, maternal fever, and vaginal colonization with Group B Streptococcus (GBS), including GBS bacteriuria– - have been strongly associated with neonatal infections. (29–34) Additionally, infant-related risk factors such as prematurity, congenital anomalies, low birth weight, and low Apgar scores (≤6 at 5 minutes) continue to increase susceptibility to NS (35)

The global burden of infections caused by MDR Gram-negative bacteria in neonates has grown significantly over the past two decades, especially in low-resource settings [36- 39). The primary driver of this resistance is the production of β-lactamases, which hydrolyze β-lactam antibiotics, rendering them ineffective. Molecular characterization in this study revealed a high prevalence of extended-spectrum β-lactamase (ESBL)-encoding genes: CTX-M1 (98.1%), TEM (79.2%), and SHV (33.9%). CTX-M1, being one of the most widely disseminated ESBL genes globally, has been documented as the predominant variant in sub-Saharan Africa.

The co-expression of multiple β-lactamase genes within the same isolate further underscores the complexity of resistance mechanisms and the growing trend observed in contemporary studies (37– 39]. Despite the widespread presence of ESBL-mediated resistance, many healthcare facilities in both rural and urban parts of Lagos State lacked routine screening capabilities for ESBL production at the time of sample collection. This diagnostic gap may have exacerbated the threat posed by MDR pathogens thus compromising effective clinical management.

In light of the above, we adopted molecular docking to evaluate the binding interactions between widely used antibiotics and two principal bacterial resistance targets - DNA gyrase and β-lactamase - offering valuable insights into drug efficacy and mechanisms of resistance.

DNA gyrase is a major target for fluoroquinolones. The docking simulations indicated Piperacillin (- 7.9 kcal/mol) and Ciprofloxacin (-7.8 kcal/mol) as exhibiting the strongest binding affinities to DNA gyrase. Piperacillin interacted with critical residues such as Asn46, Val43, Ala47, Val71, and Ile94 via hydrogen bonds and hydrophobic contacts, suggesting stable binding.

Ciprofloxacin, consistent with its known mechanism, formed hydrogen bonds with Gly77 and Thr165 and hydrophobic interactions with Ile78, effectively impeding DNA replication. Amoxicillin, although primarily a cell wall synthesis inhibitor, also showed notable affinity for DNA gyrase through interactions with Asp73, Gly77, Val97, and Thr165, indicating potential off-target effects. Imipenem, despite forming hydrogen bonds with Glu50 and Arg76, had a lower binding energy (-7.0 kcal/mol), implying weaker inhibition. Cefepime had the lowest affinity (-6.9 kcal/mol), correlating with its reduced action on DNA gyrase.

Beta-lactamase docking revealed that Ciprofloxacin also had the highest binding affinity (-8.2 kcal/mol), despite not being a conventional β-lactam substrate. Its interactions with residues Ala217, Ile221, and Ile279 suggest possible indirect inhibition or allosteric modulation. Piperacillin, a β- lactam, showed significant binding (-7.8 kcal/mol) through hydrogen bonds with Gln277 and Ala280 and hydrophobic interactions with Ala217 and Leu220, aligning with its known susceptibility to enzymatic hydrolysis. Amoxicillin (-7.6 kcal/mol) bound moderately via hydrophobic interactions but lacked hydrogen bonds, which may explain its poor efficacy against beta-lactamase-producing strains. Meropenem had the lowest affinity (-6.9 kcal/mol), forming minimal hydrogen bonds, a feature that underpins its relative resistance to enzymatic degradation and supports its use against resistant pathogens.

These molecular docking outcomes are consistent with phenotypic resistance patterns and the detected presence of bla-TEM, bla-SHV, and bla-CTX genes in clinical isolates, which explain the poor efficacy of antibiotics such as ampicillin, gentamicin, and cefotaxime. The strong affinity of Piperacillin and Amoxicillin for beta-lactamase reinforces their susceptibility to enzymatic inactivation. Conversely, Meropenem’s low beta-lactamase affinity and moderate DNA gyrase interaction indicate a dual advantage– resistance to degradation and effective target engagement.

This study underscores the value of molecular docking in predicting antibiotic effectiveness, highlighting Although Meropenem remains effective, its judicious use is imperative to prevent the rise of carbapenem-resistant Enterobacteriaceae (CRE).

## CONCLUSION AND RECOMMENDATION

Several major findings came from this research. In EOS and LOS, Klebsiella pneumoniae and Coagulase-negative Staphylococci predominated. An enhanced conventional identification method revealed two previously unreported neonatal sepsis pathogens, Enterobacter gergoviae and Serratia rubidiae. However, sequencing, BLAST alignment, and docking revealed different sequence identities and corroborated their novelty in NS in the research population.

Preventing neonatal sepsis is important, but early identification and focused antibiotic therapy are crucial. This study found severe diagnostic accuracy limitations. Blood culture is the gold standard for diagnosis, but this study emphasizes the need for molecular identification of antimicrobial resistance genes to guide therapy, especially as empirical treatments like ampicillin, gentamicin, and cefotaxime become ineffective due to rising resistance rates. Meropenem was the most effective antibiotic against resistant bacteria in this cohort.

Sequencing-based molecular diagnostics are outperforming older confirmatory approaches, which have disadvantages. Gene sequencing is useful for optimising treatment and reducing NS-related morbidity and mortality. In this work, phenotypic and molecular methods improved newborn sepsis pathogen detection and characterisation. Enterobacter gergoviae and Serratia rubidiae, previously unreported, highlight the necessity of thorough diagnostics in revealing shifting pathogenic ecosystems in Lagos healthcare settings.

More importantly, molecular docking helps understand resistance processes and select antibiotics. Molecular docking analyses revealed potential antibiotic-target interactions, supporting these findings. It explains how resistance pathways reduce neonatal sepsis medication efficacy and helps genetic and structural investigations in therapeutic options. An integrated neonatal care system requires strict antibiotic-use restrictions, regional bacterial surveillance, and resistance monitoring.

Overall, this study recommends routine ESBL gene screening in neonatal care facilities in Lagos State, given the high prevalence of multidrug resistance (MDR). Gene sequencing and molecular docking analyses should be used alongside the Microbact™ identification system, previously considered the definitive diagnostic tool for NS. Meropenem’s clinical value was strengthened by molecular docking and gene sequencing data that accurately assessed antibiotic efficacy. These sophisticated methods promise better diagnosis and patient management.

Consequently, this approach advances genotype-informed treatment and strengthens antibiotic stewardship in regions with rising antimicrobial resistance, guiding empirical treatment strategies and region-specific antibiotic stewardship initiatives to control neonatal infections. Thus, meropenem can treat NS for a long time when used wisely.

## Abbreviations

AMR: Antimicrobial resistance
CLSI: Clinical and Laboratory Standards Institute
EOS: Early-onset sepsis (≤ 28 days postpartum)
ESBL: Extended Spectrum Beta-Lactamase
GBS: Group B streptococcus
MDR: Multi-Drug resistance
NS: Neonatal Sepsis
Pi: Protein Isoelectric point

## AUTHORŚ CONTRIBUTIONS

AM Deji-Agboola, BA Iwalokun, and TA Banjo were involved in the following roles: conceptualization, development, analysis, and critical review; article manuscript draughting; sequencing, molecular docking, editing, study interpretation, and project management.

Research strategy, technical lab work, sequencing, analysis, result interpretation, review, and editing were all areas in which HO Adegboyega had a hand. PB Fakunle, OM Fatungase, MM Adeyanju, and RA Akindele helped with the analysis, interpretation, and editing of the paper. FA Oladoja helped with the analysis, interpreted the results, and prepared the manuscript, which included writing the first draft, reviewing it, editing it, and even doing molecular docking.

WB Mutiu, OA Osinupebi, TO Shorunmu, BO Ajayi, O. Onadeko, AAA Oyelekan, SO Ariyibi, and ES Omirin were all involved in the study’s execution, review and analysis of laboratory procedures and outcomes. KS Oritogun contributed to the manuscript’s development, review, editing, and statistical analysis. All authors assessed and approved the final manuscript for publication.

## DATA AVAILABILITY

All data generated or analyzed during this study are included in this manuscript.

## DECLARATIONS

### Ethical considerations

Ethical approval and authorisation were secured from the research ethics committee of the Nigerian Institute of Medical Research. Written informed consent was acquired from the mothers of the neonates after clarifying the study’s purpose and benefits, ensuring the utmost confidentiality of their information throughout the research.

### Clinical trial number

‘Clinical trial number: not applicable.’

### Consent for publication

Not applicable.

## REQUEST FOR APC WAIVER

All the authors passionately request for consideration for full waivers for the full Article Processing Charge (100% discount of the APC) as we are based in low-income country, Nigeria.

